# Nutritional Status and its Related Factors in Khalwa Residents, Khartoum state, Sudan

**DOI:** 10.1101/610972

**Authors:** Israa Hassan Giha, Mahasin Ibrahim Shaddad, Abdulrahman Yusuf, Ibrahim Abba Paga, Mounkaila Noma, Mamoun Homeida

## Abstract

**Introduction:** Khalwa is an Islamic educational institution, known as such in Sudan and called elsewhere Koranic institution. Our research aimed to assess the nutritional status and its related factors among Khalwa residents in Khartoum State.

**Methods:** A facility-based cross-sectional study was implemented in two localities of Khartoum State. A multistage sampling technique was used to selected 1273 residents. At first level, four khalwa were selected in the seven localities of Khartoum State through a stratified random sampling technique. At second level, in each of the khalwa selected, all the residents fulfilling the inclusion and exclusion criteria were included in the study. The collected data were firstly summarized numerically and graphically. Then, associations/differences among variables were determined through chi-square tests and ANOVA. A multinomial logistic regression established the relationship between the nutritional status of the residents and its related factors. All statistical tests were considered statistically significant when *p*< 0.05.

**Results:** The age of 1273 residents varied from 6 to 60 years with an average age of 15 years. Their mean body mass index (BMI) of 16.6±3.4 ranged from 7.8 to 34.0. 73.8% (939/1272) of the residents were undernourished, 23.9% (23.9%, 309/1272) were well nourished and 2.3% (29/1272) were overweight/obese. The statistically significant factors related to the nutritional status of the residents were age (under-nourished *p*=0.000; well-nourished *p=*0.004), status in the khalwa (*p*=0.001 vs *p*=0.075), resting time (*p*=0.002 vs *p*=0.038), practices of hand washing (*p*=0.165 vs *p*=0.011) and exercising (*p*=0.032 vs *p*=0.027). The food practices, despite their contributions to the model, were not statistically significant (p > 0.05).

**Discussion:** The nutritional status of khalwa should be translated urgently in a community-directed intervention based on a partnership involving the affected communities, political and administrative authorities, national, bilateral and international donors to overcome the burden of malnutrition.

## Introduction

Malnutrition challenges the world with one in three people directly affected by underweight, vitamin and mineral deficiency, overweight, obesity and diet-related non-communicable diseases. These conditions increasingly coexist in a nation, community, household, or even in the same individual across the life course. While more than 1.9 billion adults were overweight or obese worldwide in 2015, 462 million were underweight. In 2016, 155 million children < 5 years were affected by stunting while 41 million were overweight and 52 million were affected by wasting. The Global Nutrition Report Group in targeting to end malnutrition by 2030 pointed out that at least 12 out of the 17 sustainable development goals (SDGs) contain indicators that are highly relevant to nutrition; it pointed out the central role of nutrition in sustainable development [1-3]. Accurate planning and implementation of intervention programmes for reducing morbidity and mortality related to under nutrition is crucial for assessment of community nutritional status. WHO Global Database on Child Growth and Malnutrition recommended the use of anthropometric measurements as biomarkers to reflect the overall nutritional status of a human being and to classify the spectrum of different nutritional problems [4,5]. Over 40 % of the global population suffer from water scarcity and at least 663 million people worldwide lack access to safe drinking water. One third of the world population lack access to an improved sanitation leading to 13% practicing open defecation; only 19% of people worldwide wash their hands after potential contact with excreta. Sub-Saharan Africa and South Asia remain the regions of the world with the lowest sanitation coverage [6-8]. In 2010, according to the Human Rights Watch there was about 5 million children attending thousands of Quranic boarding schools across the globe [9]. The denomination of Quranic schools and attendees varies across continents and countries. In Bangladesh, India, Pakistan, Singapour, and Somalia, the Quranic schools are named madrasah/madarassa, called tsangaya in Nigeria, Daaras in Mauritania and Senegal; in Egypt the koranic schools are known as alkotab, they are commonly named khalwa in Sudan [10-14]. Khalwa is a historical Islamic educational institution in Sudan, which despite its contribution to the Islamic education has experienced little change over the years [10,11]. It mostly depends on Islamic endowments, individual donations and charities; the quality of life of the residents is impacted by poor environmental and sanitation conditions associated to an unsatisfactory diet. It remains the main center of educational opportunity for many remote rural Sudanese people [15, 16].

Various researchers were interested in the nutritional status in institutions because of its direct health impact, particularly in Quranic schools. A descriptive cross-sectional study assessed the socioeconomic, demographic and health problems on a sample of 377 Almajri of Sokoto State in Northern Nigeria. The authors reported that 68.4% of the study participants belonged to poor socio economic families, 62.8% came from polygamous families, 59.7% had urinary tract infection and 17.8% suffered from skin disease. 60.4% took bath once a month and 66.7% did not practice tooth brushing [17]. In the same country, a cross-sectional study on a sample of 360 almajiri evaluated the nutritional status and prevalence of intestinal schistosomiasis in Kawo District, Kaduna Metropolis. The results pointed out that severe malnutrition (body mass index < 15.3) prevailed more in infected children (n=67) than in non-infected (n=293) with a prevalence of respectively 75.0% and 67.0%. These findings were recently confirmed by Mohamed A et al. in East Nile locality, Sudan when assessing health and biosocial aspects of children of Quranic schools. Their results revealed that students harboring schistosomiasis and 34.5% of the students suffered from stunted growth while 17.0% were malnourished [18,19]. Further health problems in Quranic schools were reported through a comparative cross-sectional study on a sample of 213 Almajiris and 200 public school pupils seeking to evaluate the prevalence and correlates of psychiatric disorders. The study revealed a mean number of traumatic events significantly higher (1.38±1.05) among Almajiris compared to public school pupils (0.87± 0.83) with *p-value*< 0.001. Depression, enuresis, substance use remained significantly higher among Almajiris; while separation anxiety was significantly experienced by public school children [20].

Nutritional status is multifactorial involving both nutritional, biological, environmental, psychological and physical factors. A descriptive cross-sectional study on a sample of 384 students of four primary education institutions (Khalwa) assessed water and environmental sanitation services of primary education at East Nile locality (Khartoum, Sudan). The findings revealed that 75% of the Khalwa had just two meals/day, regarding the main source of safe drinking water, wells remained the first source of water for 50% of the Khalwa (n=4). For waste management half of the Khalwa did not have containers for storing solid waste which was consequently burned. Despite, 96.4% of the students were educated solely in Khalwa, only 2/3 (67.2%) of them knew about the importance of personal hygiene [21]. An analytical cross-sectional study carried out in Gezira State, Sudan on a sample of 180 students of Quranic school, revealed a prevalence of anemia of 88.33%, The prevalence was higher (47.11%) in children aged 11-14 years than in those of 7-10 years (38.15%) and 15-18 years (3.07%). The majority of the study participants had iron deficiency anemia, followed by haemolytic, macrocytic and sickle cell anemia [22]. Kheir.A.E.M. et al., in a cross-sectional survey on a sample of 406 males residing in a traditional Quranic school, revealed a prevalence of night blindness of 24.0%, conjunctival xerosis was 12.5%; and Bitot’s spots was 1.0%. A statistically significant association (*p*=0.023) was found between the duration of stay (cut-off of 6 months continuously) at the institute and the development of night blindness [23].

Khalwa, a dedicating place for teaching and memorization Quran and hadith, is a close environment where live students, teachers, supervisors under the overall responsibility of al faki or sheikh. Our research aimed to assess the nutritional status of the residents and its associated factors in four Khalwa, distributed in two of the seven localities of Khartoum State, Sudan.

## Materials and Methods

A facility-based cross-sectional study was implemented in two (Karari and Shareq Alneel) of the seven localities of Khartoum State comprising an estimated total population of 7,993,851 people in 2018 per Sudan Census Bureau of Statistics (http://www.cbs.gov.sd/). A multistage sampling technique was used to select the study participants. At first level a stratified random sampling technique was used to select the Khalwa proportionally to size of each locality based on the number of residents. At second level, two Khalwa were randomly selected in each of the two localities included in the study. At third level, in the selected Khalwa all the residents who fulfilled the inclusion and exclusion criteria (living in daily basis in selected Khalwa) and who provided their well verbal informed consent were included in the research.

The data were collected through a standardized interviewer-administrated tool, firstly developed in English and translated in Arabic for easy understanding by the study participants. The questionnaire was pre-tested prior to the study implementation in Al Khartoum and Ombadda localities.

The questionnaire comprised two parts. Part, 1 related to the institution, enabled to collected data, from each Khalwa through observations and interviews to gather statistics on residents, accommodation, sources of water and energy as well as environmental hygiene and sanitation including waste management. Part 2 recorded data related to the study participants through four subheadings the sociodemographic characteristics of the participants, their mode of living, their nutritional practices and their anthropometric measures (height, weight and Body mass index). The anthropometric measures weight (kg) and height (cm) were recorded for all the residents in using Floor type weight scale (model Zt-120). The weight and height were measured in having each resident in minimum clothes, without shoes and having each the head upright and looking straight forward. A relational database was developed through EPI-Info™ 7.1.5.2 2 for easy data entry, to avoid duplication and to minimize data entry errors in the two parts of questionnaire linked through a unique code created based of the Khalwa code and the participant identification number.

The statistical package for social sciences (SPSS version 23) was used to summarize the data numerically (mean, standard deviation, median) and graphically (frequency tables and graphics). Chi-square tests were used to determine association between categorical variables; analysis of variance determined the statistical association between continuous and categorical variables. A multinomial logistic regression was performed to establish the relationship between the nutritional status of the residents and a set of explanatory variables including age, status, duration of living in khalwa, resting time, physical activities and nutritional practices of the residents. All statistical tests were considered as statistically significant when *p*< 0.05.

## Results

### Profile of the Khalwa Surveyed in Karari and Shareq Alneel Localities

In the overall, the four Khalwa reported at the time of the research a total of 2726 residents with an average of 54 students per trainers (teachers and supervisors). With an average of 35 residents per bedroom and 5 residents per bed, the Khalwa were crowded. General network (75%, 3/4) was the first source of water which was stored in containers and traditional coolers in all the Khalwa. Uncovered zeer without tape (2 Khalwa/4) and basin (2/4) were used in half of the Khalwa whereas the recommended water storage, covered drums zeer with tap was found in one Khalwa only. Half of the Khalwa was connected to the general network as primary source of energy, half relied on generators and solar energy was the main source of energy in one institution.

A total of 88 latrines were recorded in the four Khalwa out of which 81 were in use which represented 92.0% (81/88) in term of availability of this commodity. The average number of residents per latrine used was 34 residents; this average ranged from 20 to 26 residents/latrine in Shareq Alneel locality to 103 to 207 residents/latrine in Karari locality. In the overall, an average of 32 residents shared a bathroom, this average varied from 20 to 26 residents/bathroom in Shareq Alneel to 74 to 155 residents/bathroom in Karari. For waste management, open burned system was used by 75.0% (3/4) of the Khalwa, this system was combined with car waste collection system by two, all located in Shareq Alneel locality (table 1).

**Table 1:** Commodities and waste management system available in the four Khalwa of Karari and Shareq Alneel localities.

| Commodities | Karari |  | Shareq Alneel |  | Total | % available<sup>d</sup> |
| --- | --- | --- | --- | --- | --- | --- |
|  | Mohamed Alhabib /Mojamah Zo | Almanara / Osman Ibn Afaan | Khalifa Mohamed Zein Alabdin | Maseed Elsheikh ALobeid Wd B |  |  |
| Type of latrines |  |  |  |  |  |  |
| Improved Pit latrine | . | 1 | 1 | 1 | 3 | 75.0 |
| Pit latrine | 1 |  |  |  | 1 | 25.0 |
| Open defecation | 1 |  |  |  | 1 | 25.0 |
| Use of latrines |  |  |  |  |  |  |
| Total number of Latrines | 4 | 5 | 49 | 30 | 88 |  |
| Latrines in use | 3 | 5 | 49 | 24 | 81 | 92.0 |
| Average residents/Latrine used<sup>a</sup> | 207 | 103 | 20 | 26 | 34 |  |
| Use of bathrooms |  |  |  |  |  |  |
| Total number of bathrooms | 4 | 7 | 49 | 30 | 90 |  |
| Number of bathrooms in use | 4 | 7 | 49 | 24 | 84 | 93.3 |
| Average residents/bathroom used<sup>b</sup> | 155 | 74 | 20 | 26 | 32 |  |
| Waste Management |  |  |  |  |  |  |
| Open burned system | 1 |  | 1 | 1 | 3 | 75.0 |
| Waste car collection system | . |  | 1 | 1 | 2 | 50.0 |
| Residents<sup>c</sup> | 620 | 516 | 963 | 627 | 2726 |  |

a: computed as number of residents per khalwa/number of latrines used. b: number of residents per khalwa/number of bathroom used. c: total number of residents (teachers+supervisors+students) as reported at the time of the survey. d: Percentage calculated as total number of commodities reported/4 Khalwa x 100, or as the number of commodities used/total number of commodities x 100.

### Characteristics of the Residents

A total of 1273 residents was included in our survey. The age of the residents varied from 6 to 60 years with a median age of 15 years. They lived in their respective Khalwa for an average of 1 year ranging from 0 to 35 years; 59.7% (760/1273) were in their Khalwa for 1 to 4 years. The majority of the residents were students (94.4%, 1201/1273) and the remaining 5.6% (72/1273) were either teachers or supervisors. The mean duration of living in the Khalwa of 8.3 years±8.2 was higher for supervisors than teachers (6.3 years±6.9) and students (1.3 years±1.3) and this difference was statistically significant (*p*=0.000). The association between age of the residents and the years of living in Khalwa was also statistically significant (*p*=0.000).

### Mode of living of the study participants

#### Resting time of the residents

The resting time (in hour) of the residents were estimated by subtracting the time of waking up from the one of going to bed. The resting time of the residents ranged from 5 to 8 hours with an average of 6.7 hours±0.7. This average differed from one Khalwa to another, it was lower (6.3 hours) in Almanara/Karari and Khalifa Mohamed Zein Alabdin/Shareq Alneel compared to Mohamed Alhabib/Karari and Maseed Elsheikh/Shareq Alneel where the residents had 7 hours of sleep. According to the age of the residents, the resting time was lower in the younger age groups: 6.4 hours±0.6 (< 15 years) and 6.9 hours±0.6 (15-24 years). In the age groups ≥ 25 years it ranged from 7.1±0.8 (≥35 years) to 7.2±0.6 (25-34 years). The resting time of the students of 6.6 hours±0.6 was lower than of the teachers (7.0 hours±0.8) and supervisors (7.1 hours±0.8). A statistically significant difference was found between resting time and institutions (*p*=0.000), age groups (*p*=0.000) and status of the residents (*p*=0.000).

#### Activities of the residents

The residents of the Khalwa performed five main activities namely reading and writing Quran, walking, cooking, cleaning their respective Khalwa and washing their clothes. *Reading and writing Quran* was the predominant activity (100.0%, 1273/1273) in all the Khalwa. According to the Khalwa, *walking* is performed daily by 95.9% (1212/1273) of the residents, daily *cooking by* 59.6% (716/1202) whereas, cooking was an occasional activity for 40.3% (485/1202). *Cleaning their respective Khalwa* was an activity implemented once a week by 78.2% (995/1273) of the residents and 94.5% (1203/1273) of the participants *washed their clothes* once a week. According to the status of the residents, walking was practiced by all the teachers (n=36), supervisors (n=36) and students (n=1201). The cooking activity was performed mainly by the students (100.0%, 1201/1201) who carried the activity on a daily basis (59.6%, 716/1201) and occasionally (40.4%, 485/1201). All the three bodies participated in cleaning their Khalwa, mainly the students (91.0%, 1093/1201) followed by the supervisors (38.9%, 14/36) and the teachers (22.2%, 8/36).

#### Hygiene and Sanitation

Of the 1273 participants, 67.6% (861/1273) practiced hand washing without soap, 31.0% (394/1273) with soap, 0.8% (10/1273) reported washing hands with sand and 0.6% (8/1273) did not practice hand washing. The practice of hand washing was regrouped as “good practice” when participants washed hands with soap and all other three practices of hand washing was labeled as “poor practice”. In the overall, 69.0% (879/1273) of the residents had poor practice of hand washing and the remaining 31.0% (394/1273) had good practice. A statistically significant difference was found between the practice of hand washing across Khalwa (*p*=0.000), status of the residents (*p*=0.000) and age of the participants (*p*=0.000). All the residents across the four Khalwa used “miswak” for teeth brushing before each prayer. 23.2% (294/1267) of participants brushed once a day their teeth with paste and the remaining 76.8% (973/1267) brushed their teeth without paste. Teeth brushing with paste was statistically different (*p*=0.000) across Khalwa and the status of residents. Student had the lowest prevalence of teeth brushing (19.7%, 235/1195) with paste.

#### Nutritional practices and intakes in the Khalwa

In general, the residents had two meals a day, breakfast at 10:00 am for 96.5% (1228/1273) and at 11:00 am for the remaining 3.5% (45/1273). The second meal (lunch) was served at 5:00 pm for the majority (99.5%, 1267/1273) and at 4:00 pm for 6 residents (0.5%). 32 (2.5%) residents had dinner at 10:00 pm. Of those having dinner, 52.8% were teachers, 27.8% supervisors and 0.2% students. Almost all the residents (96.6%, 1227/1270) reported that they needed extra meals; they were students (100.0%, 1119/1119), teachers (52.8%, 19/36) and supervisors (9/35).

*Assida* (sorghum seeds crushed and cooked) and served with *mullah* (water, scanty grinded dried meat and okra) was the food consumed on a daily basis in all the Khalwa by respectively 99.6% (1261/1266) and 99.7% (1263/1267) of the residents. *Saliga/fatta* (meat soup with bread) was served either occasionally (55.2%, 703/1273) or on a weekly basis (44.5%, 566/1273). *Beans* (for preparing tamia, foul, balila, and loubia), were generally served occasionally (54.8%, 480/876) or every day (44.6%, 391/876). The frequency of consumption of bread was occasionally (95.1%, 523/550), daily (2.7%, 15/550) and weekly (12/550).

The consumption of fresh fruits was not a habit with 5 residents out of which 4 (3 teachers and 1 supervisor) had fruits weekly and a teacher daily. Dattes (dry fruits) were consumed occasionally (72.3%, 910/1258), daily (27.3%, 343/1258) and weekly (0.4%, 5/1258). Having dry fruits on a daily basis was more frequent in teacher population (42.9%, 15/35) than in supervisors (40.0%, 14/35) and students (26.4%, 314/1188).

*Nashah* (porridge made of cereals and sugar) was consumed by all (1273/1273) the residents on a weekly basis, *naturel juice* (hisbiscus, conceales) by 1.9% (24/1273) on a daily basis by 6 residents and occasionally by 18. 1.2% (15/1273) had *milk* every day (7 residents) and occasionally (8 residents). The majority (74.8%, 943/1261) of the residents had tea on a daily basis, 24.7% (312/1261) occasionally, 0.3% (4/1261) once a week and 0.2% (2/1261) never took tea. Coffee was reported to be drunk every day by 14.8% (67/454) of the residents and occasionally by 85.2% (387/454).

### Anthropometric measurements

The median weight of the residents (n=1272) of 36 kg ranged from 11 to 107 kg. According to the Khalwa, the weight varied from 34.3 kg±13.7 to 43.0 kg±12.6. The students had a mean weight lower (37.6 kg ±12.7) than the teachers (60.1 kg±16.6) and the supervisors (62.6 kg±24.5). A statistically significant difference (*p*=0.000) was found between the weight and the status of the residents. With regard to the duration of living in Khalwa, the mean weight was lower (37.2 kg±13.0) for those residing for 1-4 years than those living for < 1 year (38.8 kg±13.2) and ≥ 5 years (61.2 kg±19.0). A statistically significant difference was found between the age of the participants, the duration of living in Khalwa and the weight with a *p-value* of respectively 0.000.

The 1272 residents had a height ranging from 1.1 m to 1.9 m with a median height of 1.5 meter. According to their respective Khalwa, the mean height of the residents (n=1272) of 1.5 m±0.2 varied between 1.4 m±0.1 and 1.5 m±0.2. The lowest mean height (1.5 m±0.2) was recorded in students. The mean height increased with the age of the residents and their respective length of living in their respective Khalwa. A statistically significant difference (p=0.000) was found between Khalwa, status of the residents, age of the study participants, length of living and height of the residents.

The mean body mass index (BMI) of the residents (n=1272) of 16.6±3.4 ranged from 7.8 to 34.0. This mean BMI varied across Khalwa, status and age of the residents with a statistically significant difference (*p*=0.000). The cutoff points proposed by the World Health Organization were used for classification of the nutritional status of the residents. The majority (73.8%, 939/1272) of the residents were underweight (BMI < 18.5 kg/m^2^), 23.9% (304/1272) had normal weight (BMI 18.5-24.9 kg/m^2^), 2.0%, (25/1272) were overweight (BMI ≥ 25 kg/m^2^) and 0.3% (0.3%, 4/1272) were obese (BMI ≥ 25 kg/m^2^).

According to WHO classification, the BMI of the residents was regrouped in three categories namely undernourished (underweight), well nourished (those with normal weight) and overweight/obesity. 73.8% (939/1272) of the residents were undernourished, 23.9% (23.9%, 304/1272) were well nourished and 2.3% (29/1272) were overweight/obese. The prevalence of undernourished of 73.8% ranged from 65.9% in Karari Mohamed Alhabib to 86.2% in Shareq Alneel Khalifa Mohamed Zein Alabdin as revealed by table 2 This average prevalence ranged from 30.6% (11/36) among the supervisors to 76.4% (917/1201) among the students. The prevalence of undernourished was higher 93.5% (589/630) in the age group < 15 years. According to the duration of living in Khalwa, the prevalence of undernourished was higher (76.8%, 584/760) in those living since 1-4 years in Khalwa compared to the residents living < 1 year (73.8%, 333/451) and ≥ 5 years (36.1%, 22/61).

**Table 2:** Nutritional status of the residents according to their Khalwa, status, age and duration of living in their respective institution (n=1273)

| Variable | Nutritional status |  |  |  | % under-nourished |
| --- | --- | --- | --- | --- | --- |
|  | Undernourished | Well nourished | Overweight/Obesity | Total |  |
| Khalwa |  |  |  |  |  |
| Shareq Alneel Khalifa Mohamed Zein Alabdin | 193 | 23 | 8 | 224 | 86.2 |
| Karari Almanara | 346 | 86 | 10 | 442 | 78.3 |
| Shareq Alneel Maseed Elsheikh | 187 | 90 | 6 | 283 | 66.1 |
| Karari Mohamed Alhabib | 213 | 105 | 5 | 323 | 65.9 |
| Total | 939 | 304 | 29 | 1272 | 73.8 |
| Status of the residents |  |  |  |  |  |
| Teacher | 11 | 15 | 9 | 35 | 31.4 |
| Supervisor | 11 | 12 | 13 | 36 | 30.6 |
| Student | 917 | 277 | 7 | 1201 | 76.4 |
| Total | 939 | 304 | 29 | 1272 | 73.8 |
| Age group of the residents |  |  |  |  |  |
| < 15 years | 589 | 41 | 0 | 630 | 93.5 |
| 15-24 years | 329 | 230 | 7 | 566 | 58.1 |
| 25-34 years | 9 | 23 | 8 | 40 | 22.5 |
| ≥ 35 years | 4 | 10 | 14 | 28 | 14.3 |
| Total | 931 | 304 | 29 | 1264 | 73.7 |
| Duration of living in Khalwa |  |  |  |  |  |
| < 1 year | 333 | 115 | 3 | 451 | 73.8 |
| 1-4 years | 584 | 168 | 8 | 760 | 76.8 |
| ≥ 5 years | 22 | 21 | 18 | 61 | 36.1 |
| Total | 939 | 304 | 29 | 1272 | 73.8 |

### Relationship between the nutritional status of the Khalwa residents and their nutritional practices and environmental factors

A multinomial logistic regression was performed to assess the relationship between nine explanatory variables and the nutritional status of the residents. This later was prevalent as under-nourished (73.8%, 939/1272), well-nourished (23.9%, 304/1272) and overweight/obesity (2.3%, 29/1272).The nine predictors used were namely the age of the residents in years, the duration of living in Khalwa in years, the status of the residents (teacher, supervisor, student), the resting time (sleep), hand washing (good and poor practice), exercising daily (yes=every day; no=once a week/occasionally), food intake measured as food2 (meals made of beans) and food4 (yes= having once a week Saliga/Fatta; no=occasionally/never) and dry fruit consumption (yes=having dattes every day; no=dattes served once a week/occasionally).The reference subpopulation for nutritional status used in the multinomial regression model was “overweight/obesity” which was compared with the subpopulations “under-nourished” and “well-nourished” for each of the predictive factors.

#### Under-nourished relative overweight/obesity

Age (*p*=0.000), status of the residents (*p*=0.0001), resting time/sleep(p=0.002) and exercising (*p*=0.032) were statistically associated to the nutritional status of the residents. Having daily dattes despite not statistically significant (*p*=0.056) contributed to explain the nutritional status of the residents by 4.4 times [95% CI:0.965-20.316]. Eating salitiga/fatta weekly contributed to the nutritional status by 2.8 times [OR=2.804, 95% CI:0.12-65.265] despite a no statistically significant association (*p*=0.521). Hand washing practice [OR= 0.369, 95% CI:0.09-1.508, *p*=0.165] with a coefficient of −0.996 indicated that the practice of hand washing was more in under-nourished resident students. Having meals made with beans [OR= 0.246, 95% CI:0.031-1.964, *p*=0.186] had a negative contribution to the model with a coefficient of −1.402 indicating eating meals processed with beans occasionally/every week led more likely to be undernourished (table 3).

**Table 3:** Multinomial regression model predicting the nutritional status in Khalwa based on the age, status, duration of living in Khalwa, hygiene and dietary habits when using overweight/obesity as reference subpopulation.

| Subpopulation | Variable | B | Std. Error | Wald | df | p | OR | 95% CI OR |  |
| --- | --- | --- | --- | --- | --- | --- | --- | --- | --- |
|  |  |  |  |  |  |  |  | Lower | Upper |
| Under-nourished | Intercept | 15.072 | 5.549 | 7.377 | 1 | 0.007 |  |  |  |
|  | Age | -0.251 | 0.05 | 25.441 | 1 | 0.000 | 0.778 | 0.705 | 0.858 |
|  | Live | 0.071 | 0.069 | 1.061 | 1 | 0.303 | 1.074 | 0.938 | 1.23 |
|  | Status | 2.028 | 0.607 | 11.159 | 1 | 0.001 | 7.602 | 2.313 | 24.994 |
|  | Sleep | -1.786 | 0.581 | 9.467 | 1 | 0.002 | 0.168 | 0.054 | 0.523 |
|  | Hand washing practice | -0.996 | 0.718 | 1.926 | 1 | 0.165 | 0.369 | 0.09 | 1.508 |
|  | Exercising | -1.045 | 0.487 | 4.606 | 1 | 0.032 | 0.352 | 0.135 | 0.913 |
|  | Food2 (made with beans) | -1.402 | 1.06 | 1.75 | 1 | 0.186 | 0.246 | 0.031 | 1.964 |
|  | Food4 (Saliga/Fatta once a week) | 1.031 | 1.606 | 0.412 | 1 | 0.521 | 2.804 | 0.12 | 65.265 |
|  | DryFruits1 (having dattes every day) | 1.488 | 0.777 | 3.664 | 1 | 0.056 | 4.428 | 0.965 | 20.316 |
| Well- nourished | Intercept | 15.233 | 5.339 | 8.14 | 1 | 0.004 |  |  |  |
|  | Age | -0.104 | 0.036 | 8.499 | 1 | 0.004 | 0.901 | 0.84 | 0.966 |
|  | Live | -0.051 | 0.056 | 0.848 | 1 | 0.357 | 0.95 | 0.852 | 1.059 |
|  | Status | 0.756 | 0.425 | 3.168 | 1 | 0.075 | 2.13 | 0.926 | 4.9 |
|  | Sleep | -1.156 | 0.559 | 4.284 | 1 | 0.038 | 0.315 | 0.105 | 0.94 |
|  | Hand washing | -1.787 | 0.706 | 6.407 | 1 | 0.011 | 0.167 | 0.042 | 0.668 |
|  | Exercising | -1.006 | 0.454 | 4.913 | 1 | 0.027 | 0.366 | 0.15 | 0.89 |
|  | Food2 (made with beans) | -0.842 | 1.044 | 0.651 | 1 | 0.42 | 0.431 | 0.056 | 3.331 |
|  | Food4 (Saliga/Fatta once a week) | -0.393 | 1.512 | 0.068 | 1 | 0.795 | 0.675 | 0.035 | 13.074 |
|  | DryFruits1 (having dattes every day) | 0.379 | 0.732 | 0.269 | 1 | 0.604 | 1.461 | 0.348 | 6.13 |

#### Well-nourished relative overweight/obesity

Age of the residents [OR=0.901, 95% CI:0.84-0.966, *p*=0.004], resting time [OR=0.315, 95% CI:0.105-0.94, *p*=0.038], practice of hand washing [OR=0.167, 95% CI:0.042-0.668, *p*=0.011], exercising/walking every day [OR=0.366, 95% CI:0.15-0.89, *p*=0.027] had all a statistically significant association with the nutritional status of the residents. The status of the residents contributing to the model by 2.13 times [95% CI:0.926-4.9, *p*=0.075] was not statistically associated to the nutritional status; as well as having daily dattes [95% CI:0.348-6.13, *p*=0.604] contributing by 1.5 times. The duration of living in Khalwa, the meals processed with beans or eating saliga/fatta, were not statistically significant but their negative contributions indicated that as the nutritional status (well-nourished) increased, those explanatory variables decreased (table 3).

In the overall, the nutritional status of the residents of the Khalwa surveyed were statistically associated with the age of the residents (under-nourished *p*=0.000; well-nourished *p=*0.004), the status of the residents statistically significant (*p*=0.001) in under-nourished was not statistically significant (*p*=0.075) in well-nourished subpopulation. The resting time was statistically associated with the nutritional status in both under-nourished and well-nourished with a *p-value* of respectively 0.002 and 0.038. The practice of hand washing not statistically associated with the nutritional status in under-nourished (*p*=0.165) was statistically associated with the nutritional status in well-nourished subpopulation (*p*=0.011) with a negative contribution in both subpopulations of respectively −0.996 and −1.787 indicating in both subpopulations when the nutritional status improved, the practice of hand washing decreased.

A statistical significant association between nutritional status and exercising in both subpopulations was found (exercising in under-nourished *p*=0.032 vs exercising in well-nourished *p*=0.027) with a contribution to the model of respectively −1.045 and −1.006 indicating better the nutritional status, poor was the practice of exercising.

## Discussion

In our study, the residents, all males (n=1273) with an average age of 15 years (range:6-60 years) were teachers (2.8%), supervisors (2.8%) and students (94.4%). This last population had an average age of 14.6 years±3.8 (range:6-38 years). The residents had two meals/day with 96.6% in need of extra meals. This was in line with Mohamed et al. [21] who indicated that two meals were daily served in 75.0% of the khalwa surveyed and 25.0% had one meal per day. In other institutional settings, the numbers of meals served to secondary school students varied from two meals (10.5%) to more than three/daily (1.4%) with 88.1% having three meals/day [24]. Assida (99.6%) and mullah (99,7%) were the foods daily served to the residents whereas saliga/fatta, were served occasionally. Nutritional practices varied from cultures and societies. Kheir A.E.M. et al. [23] reported that in a Traditional Quran Boarding School in Sudan, boys diet consisted most of a staple cereal (sorghum or millet) and “mullah” (water, scanty, grinded dried meat and okra) whereas in Nigeria [25] 67.1% of the foods consumed were cereals; 10.1% were roots and tubers consumption, legumes were 21.1%, vegetables 16.4%, milk and products 3.1%, meat and fish 3.4% and fruits 4.1%.

Our study revealed that the resting time of the residents ranged from 5 to 8 hours with an average of 6.7 hours±0.7. This average was lower (6.6 hours±0.6) in the group of students than in teachers (7.0 hours±0.8) and supervisors (7.1 hours±0.8) with a statistically significant difference (*p*=0.00) between the status of the residents and their resting time. Musaiger.A.O. et al. [26] in their research on obesity, dietary habits, and sedentary behaviors among adolescents in Sudan reported that 28.0% (112/400) of the participants slept for < 7 hours/day and 72.0% (288/400) had ≥ 7 hours sleep. Our research did not focus on mental health which could impact the nutritional status of a community, however we took note of the findings of Abubakar-Abdullateef et al [13] who indicated in a comparative study on a sample of 213 Almajiris and 200 public school pupils that the proportion of participants having any mental health problems among the Almajiri and public school pupils was 57.7% and 37% respectively.

Regarding the physical activities, 95.9% of our study participants reported walking daily, cooking was daily performed by 59.6% predominately by students; 94.5% of the participants washed their clothes once a week and 78.2% cleaned their institution once a week. In assessing the nutritional status of secondary school adolescents (13-18 years), Abdalla.M.M.A et al [24] reported that 73.8% of their study population (n=210) practiced physical activities and 26.2% did not. The frequency of the physical activities was daily (47.6%), two-three times/week (26.2%) and never (26.2%). The types of physical activities performed were predominately football (51.0%) followed up by swimming (19.0%), jogging (2.9%) and walking (1.0%).

Regarding hygiene and sanitation, 31.0% of our study population had a good practice of hand washing, 23.2% of the residents brushed once a day their teeth with paste. The four Khalwa were crowded with an average of 35 residents/bedroom and 5 residents/bed. Improved pit latrine was the most frequent types of latrine available (75.0%,3/4) and open burned system was used by 3 of the 4 Khalwa for waste management. In their study on water and environmental sanitation services available to primary education (Khalwa), Mohamed et al [21] revealed that 98.4% of the students practiced hand washing with not all of them using soap. The benefits of brushing teeth were not known by 9.4% and 90.6% recognized the religious prescription of teeth brushing. The mean BMI of the residents (n=1272) of 16.6±3.4 ranged from 7.8 to 34.0; it varied across Khalwa, status and age of the residents with a statistically significant difference (*p*=0.000) 73.8% of the residents were undernourished, 23.9% were well nourished and 2.3% were overweight/obese. In Sudan, various authors discussed the nutritional status in different institutions and across age groups. Faroug Bakheit Mohamed Ahmed et al. [27] found that 87.2% of children had normal nutrition status and the remaining 12.8% suffered from malnutrition whereas Musaiger.A.O. et al. [28] revealed that 20.5% the study population were underweight, 14.7% were overweight and 1.7 % were obese and Ali A. A.A et al. who used anthropometry as an indicator of nutritional status of Sudanese prepubertal school children (6-9 years) [29] found that underweight was 22.34% and obesity 2.13% with no marked gender variation. Mohamed A et al. who assessed the health and biosocial aspects of children at Quranic Schools indicated that 34.0% and 17.0% of the study participants were respectively underweight and malnourished [18]. Elsewhere in the literature, Mohammed M et al. [19] assessing the nutritional status and prevalence of intestinal schistosomiasis among Al-majiri population in Kawo District of Kaduna Metropolis (Nigeria) revealed that severe malnutrition prevailed more in infected children (75.0%) than in non-infected (67.0%). Ganganahalli P et al. [30] found in a sample of 176 children of a private school that 84.1% had normal body mass index, 10.2% were overweight and 5.7% were obese. Whereas in Iran, through a screening program to assess the health and nutritional status of 2596 school children, Rezaeian S et al. [31] revealed a prevalence of wasting of 3.1%, underweight of 9.48% and stunting of 2.85%.

The limitations related to our study were its restriction to the foods practices, the hygiene and sanitation conditions of the residents, their access to safe drinking water; in the other hand, the health conditions including mental status of the residents were not investigated. However, despite those limitations, the factors related to the nutritional status of the residents were assessed through a multinomial regression analysis which revealed that age of the residents (undernourished *p*=0.000; well-nourished *p*=0.004), their status (undernourished *p*=0.001; well-nourished *p*=0.075), their resting time (undernourished *p*=0.002 vs well-nourished *p*=0.038), their practice of handwashing (undernourished *p*=0.165 vs well-nourished *p*=0.011), and their physical activity (undernourished *p*=0.032 vs well-nourished *p*=0.027) were the statistically significant contributors related to the nutritional status of Khalwa residents. The food practices despite their contributions to the model were not statistically significant (p > 0.05).

## Conclusions

Khalwa offers an opportunity of learning to low socio-economic children; furthermore, as an institution dedicating to memorizing the Quran, it was not surprising to found a high proportion of undernourished predominately in < 15 years in such close environment. This nutritional status of the residents of Khalwa should be translated urgently into proper actions to overcome the burden of malnutrition through a public-private partnership for implementing nutritional and water sanitation and hygiene (WASH) programmes alongside with education programmes.

## Acknowledgment

Our sincere thanks to Dr. Hanan Tahir, Director of Tropical and Health Programme in the University of Medical Sciences and Technology for her support and guidance throughout the research. We would like to recognize and acknowledge our study participants whose contributions were crucial to bring this research to a successful end.

## CONTRIBUTIONS OF THE AUTHORS

**IHG**: Designed and implemented the research, participated actively in the statistical analysis and read the initial drafts of the manuscript.

**MIS**: Computerized the data, participated actively in the statistical analysis and draft the manuscript.

**AY & IAP**: Assisted in data collection and coding.

**MN**: Approved the research proposal and guided the statistical analysis and participating in the proofreading.

**MH**: Participating in the proofreading/

**All the authors** read and approved the final version of the manuscript prior to its submission.

## Data Availability

In accordance to data sharing and PLOS ONE policy on the matter, the authors declared that if the submitted manuscript is accepted for publication, the data will be deposited in the generalist repository of Dryad.

## SOURCE OF FINANCING

The research was fully supported by Israa Hassan Giha in the framework of the fulfillment her Master of Sciences in Public and Tropical Health, in the University of Medical Sciences and Technology.

## CONFLICT OF INTEREST

No conflict of interest

## ETHICAL CLEARANCE

The research proposal was approved by Sumasri Institutional Review Board in the framework of community based survey. Authorization for implementation of the research was obtained from each individual Sheik of the four Khalwa surveyed. A verbal well informed consent was also obtained from each of the residents included in the research. The confidentiality was insured through the use of an anonymous research tool. The participants were reassured that the data collected from them will not be used for any other purpose other the objectives assigned to the research.

## References

1. WHO (2018). Global Nutrition Policy Review 2016 - 2017: Country progress in creating enabling policy environments for promoting healthy diets and nutrition.

2. UNICEF/ WHO/ World Bank Group. Levels and Trends in Child Malnutrition. Joint Child Malnutrition Estimates 2017 edition.

3. International Food Policy Research Institute. 2016. Global Nutrition Report 2016: From Promise to Impact: Ending Malnutrition by 2030. Washington, DC.

4. Onila, S.O, Owa J.A., Onayade A.A. & Taiwo, O. (2006). Comparative Study of Nutritional Status of Urban abd Rural Nigerian School Children. Journal of Tropical Pediatrics, 53 (1), 39–43. doi:10.1093/rropej/fmlo51.

5. WHO (1997). WHO GlobalDatabaseon Child Growthand Malnutrition.WHO/NUT/97.4.

6. United Nations. Sustainable Development Goals. Goal 6: Ensure access to water and sanitation for all. https://www.un.org/sustainabledevelopment/water-and-sanitation.

7. WHO 2015. Improving nutrition outcomes with better water, sanitation and hygiene: Practical solutions for Policy and Programmes.

8. Freeman, M. C., Stocks, M. E., Cumming, O., Jeandron, A., Higgins, J. P. T., Wolf, J., Curtis, V. (2014). Systematic review: Hygiene and health: systematic review of handwashing practices worldwide and update of health effects. Tropical Medicine & International Health, 19(8), 906–916. doi:10.1111/tmi.12339.

9. Fiasorgbor A. Dori et al. The Lived Experiences of Quranic Boarding School Pupils in the Bawku Municipality, Ghana. International Journal of Community Development, vol. 3, no. 2, 2015, 79–90. doi: 10.11634/233028791503727.

10. UNESCO. Child Family Community. ‘Koranic’ Schools in Sudan as a resource for UPEL: Results of a study on Khalwas in Rahad Agricultural Project. N.S. 160.

11. Osman Mohammad Eid. The Khalwa as an Islamic Educational Institution in The Sudan [Thesis]. University of Edinburgh, November 1985, pages 657.

12. Human Rights Watch April 2010. “Off the Backs of the Children”. Forced Begging and Other Abuses against Talibés in Senegal.

13. Abubakar-Abdullateef et al. A comparative study of the prevalence and correlates of psychiatric disorders in Almajiris and public primary school pupils in Zaria, Northwest Nigeria. Child Adolesc Psychiatry Ment Health (2017) 11:29. DOI 10.1186/s13034-017-0166-3.

14. Ishtiyaque M. et al. Role of madarassa in promoting education and socio-economic development in Mewat district state of Haryana, India. Procedia - Social and Behavioral Sciences 120 (2014) 84–89.

15. James Pruess. The “Koran” School, The “Western” School, And The Transmission of Religious Knowledge: A Comparison From The Sudan. Northeast African Studies, vol. 5, No. 2 (1983), pp. 5–39.

16. Patrick D. Lynch Abasalih Al-Fatih Qarib Allah Saifelislam M. Omer, (1992), “Educational Change and the Khalwa in the Sudan: Reform Reformed”, Journal of Educational Administration, Vol. 30 Iss 4 pp. http://dx.doi.org/10.1108/09578239210020480P.

17. Shuaib H.A, Jimoh. A.O. Assessment of Socioeconomic, Demographic and Health Problems of Almajiri in Sokoto state, North Western Nigeria, International journal of tropical medicine 6 (3); 58–60, 2011 doi:10.3923/ijtmed.2011.58.60.

18. Mohamed A, Mutamad A., M. Homeida et al. Health and Biosocial Aspects of Children at Quran Schools, East Nile Locality, Khartoum, Sudan. IJCMPR, 2017, pages 2386–2388.

19. Mohammed M, Vantsawa PA, Abdullahi UY, Muktar MD (2015) Nutritional Status and Prevalence of Intestinal Schistosomiasis among Al-majiri Population in Kawo District of Kaduna Metropolis, Kaduna State-Nigeria. J Bacteriol Parasitol 6: 237. doi:10.4172/2155-9597.1000237.

20. Abubakar-Abdullateef et al. A comparative study of the prevalence and correlates of psychiatric disorders in Almajiris and public primary school pupils in Zaria, Northwest Nigeria Child Adolesc Psychiatry Ment Health (2017) 11:29DOI 10.1186/s13034-017-0166-3.

21. Mohamed et al. Assessment of Water and Environmental Sanitation Services of Primary Education (Kwala) at East Nile Locality, Sudan, 2017. Internal Journal of Research -GRANTHAALYAH, Vol 6 (Iss.7): July 2018, page:16–22. doi:10.5281/zenodo.1322991.

22. Eltayeb. E.S.M. et al. Prevalence of anaemia among Quranic school (Khalawi) students (Heiran)in Wad El Magboul village, rural Rufaa, Gezira State, Central Sudan: a cross sectional study. PanAfrican Medical Journal. 2016; 24:244 doi:10.11604/pamj.2016.24.244.8355.

23. Kheir .A.E.M. et al. Xerophthalmia in a Traditional Quaran Boarding School in Sudan. Middle East Afr J Ophthalmo 2012 Apr-Jun 19 (2) 190–193. doi: 10.4103/0974-9233.95247.

24. Abdalla. M.M.A et al. Assessment of Nutritional Status of the Adolescents (13-18yrs) Studying in Secondary Schools In Elhafeir Area-Dangle Locality-Northern State 2018. Indian Journal of Applied Research. Volume-8, Issue-5, May-2018. ISSN - 2249-555X. IF: 5.397. IC Value: 86.18.

25. Ibrahim H. Assessment of Nutritional Status of two selected Tsangaya in Katsina Metropolis [Thesis]. Umaru Musa Yar’Adua University, January 2018, Katsina, Nigeria.

26. Musaiger. A.O. et al. Obesity, Dietary Habits, and Sedentary Behaviors Among Adolescents in Sudan: Alarming Risk Factors for Chronic Diseases in a Poor Country. Food and Nutrition Bulletin 2016, Vol. 37(1) 65–7.

27. Faroug Bakheit Mohamed Ahmed, Esam-Eddin Bakheit Mohamed Ahmed. Malnutrition Among Basic Schools’ Children of Elshagalwa Village, Shendi Locality, Sudan. International Journal of Nutrition and Food Sciences. vol. 5, no. 2, 2016, pp. 134–138. doi:10.11648/j.ijnfs.20160502.17.

28. Musaiger. A.O. et al. Obesity, unhealthy dietary habits and sedentary behaviors among university students in Sudan: growing risks for chronic diseases in a poor country. Environ Health Prev Med (2016) 21:224–230. DOI 10.1007/s12199-016-0515-5.

29. Ali A.A.A, Abdalla A. A., Zaki A. Z. M. Anthropometry as an Indicator of Nutritional Status in Sudanese Prepubertal School Children (6-9 years) At Alfitaih Village, Khartoum, Sudan, September 2014. World Journal of Pharmacy And Pharmaceutical Sciences, Volume 4, Issue 06, 182–191.

30. Ganganahalli P, Tondare MB, Durgawale PM. Nutritional Assessment of Private Primary School Children in Western Maharashtra: A Cross-sectional Study. Ntl J Community Med 2016; 7(2):97–100.

31. Rezaeian S et al. Assessment of Health and Nutritional Status in Children Based on School Screening Programs. Health Scope. 2014 Winter; 3(1): e14462. doi: 10.17795/jhealthscope-4462.

